# Case-Base-Control designs

**DOI:** 10.1101/723452

**Authors:** Najla Saad Elhezzani, Wicher Bergsma, Mike Weale

## Abstract

Most genome-wide association studies (GWASs) use randomly selected samples from the population (hereafter bases) as the control set. This approach is successful when the trait of interest is rare; otherwise, a loss in the statistical power to detect disease-associated variants is expected. To address this, a proposal to combine the three sample types, cases, controls and bases is introduced, for instances when the disease under study is prevalent. This is done by modelling the bases as a mixture of multinomial logistic functions of cases and controls, according to the disease prevalence. The maximum likelihood method is used to estimate the underlying parameters using the EM algorithm. Three classical tests of association; score, Walds, and likelihood ratio tests are derived and their power of detecting genetic associations under different designs is compared. Simulations show that combining the three samples can increase the power to detect disease-associated variants, though a very large base sample set can compensate for the lack of controls.

## 1 Introduction

In a typical case-control study which requires controls to be free of the disease of interest, the process of selecting and genotyping a control set for different diseases is expensive and time consuming. This led many researchers to take some expedient approaches to overcome this problem. For example, one of the aspects that characterizes most large GWASs e.g., The Wellcome Trust Case Control Consortium (WTCCC, http://www.wtccc.org.uk) [1], is the random selection of their control sets. In other words, controls are random samples from the population of interest with unknown disease status, rather than individuals who have been screened to ensure that they do not have the disease under study. The main advantage of adopting this approach is that genome-wide genetic data for various reference sample sets are freely available. Examples include the 1958 British Birth Cohort and the three national UK Blood Services, and both have been used in the WTCCC. Although this approach allows large time and cost savings to be made relative to sourcing and genotyping a bespoke control set, its results are reliable only if the disease of interest is rare, otherwise the degree of case contamination in the base set will be high. Examples include depression and sub-clinical conditions such as mild acne. In such situations, experimenters may be faced with a choice between analysis using a small control sample set or a large base sample set.

In this paper we combine three data types; controls, bases and cases, in a single design called the case-base-control design (CBC). We use “reverse regression” techniques, which treat the SNP’s genotype as the outcome and the phenotype as the predictor in a linear model. Reversed approaches have been previously used over classical approaches, for their capacity to simultaneously incorporate both quantitative and categorical phenotypes in the same model. See for example, the MultiPhen package [2] and the SCOPA software [3], which both target multiple-phenotype associations. Although “reverse regression” offers technical flexibilities, its estimates, especially those of the odds ratios are non-univocal, in the sense that they cannot be interpreted in terms of the effect of a SNP on a phenotype [3].

The “reverse regression” approach used herein, is based on the multinomial logistic regression [4] that models genotypes given disease status (case or control). Respective probabilities are then mixed according to the disease prevalence to form the base set. We investigate the effect of capturing information across all three data types in a joint analysis on the statistical power to detect disease associated variants. It is worth noting that, during our work on the CBC model, a similar idea was published by McCarthy et al. [5] to increase power to detect HIV associated variants, using cases, bases and a set of individuals demonstrating an exceptional resistance to HIV infection (HIV-1 high-risk seronegative individuals, HRSN). Although the motivation to simultaneously analyse the three groups is shared between the work of McCarthy et al. and ours, the corresponding approach and the underlying statistical problem are different. Basically, unlike our approach, McCarthy et al., prospectively modeled the probability of an individual being HRSN (*i* = 2), HIV positive (*i* = 1), or base (*i* = 0) conditional on genotype value g (reference allele count) using a *logit* model that treats the bases as the baseline category, as follows:

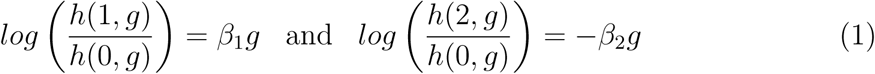

An alternative hypothesis of this form *H*_1_: at least one *β* ≠ 0 would reflect some biologically implausible genetic effect across some groups. For example, if only *β*_1_ > 0 while *β*_2_ < 0, this means having more copies of the mutant allele will increase both the probability that an individual is HIV+ (relative to bases) and the probability that an individual is HRSN (relative to bases). Such irrelevant patterns were excluded using a constrained alternative hypothesis that only reflect plausible risk or protective effects that is *H*_1_ : *β*_1_*β*_2_ > 0 with at least one *β* ≠ 0. Such formulation can capture (1) risk effect that is when *β*_1_ and *β*_2_ are both positive, in other words, having more copies of the mutant allele will increase the probability that an individual is HIV+ (relative to bases) and decrease the probability that an individual is HRSN. (2) protective effect that is when *β*_1_ and *β*_2_ are both negative. In other words when having more copies of the mutant allele will decrease the probability that an individual is HIV positive (relative to bases) and increase the probability that an individual is HRSN. The above constraints are irrelevant in our model as it is based on reverse regression techniques that assumes multiplicative genetic trends.

The outline of this paper is as follows. First, assuming known prevalence, we show the model derivation and the score test (ST) of association. For the latter, we focus on the most commonly seen trend of genetic effects on phenotypes, which is additive on the *logit* risk scale. Next, we show how the derived ST can be seen as a contrast test of the genotypes means in different groups. The resulted formulation of the ST facilitates the derivation of the power and the sample size required to achieve a specified statistical power. Further, we study the asymptotic distribution of the ST statistic under both hypotheses, and we check its power sensitivity to prevalence misspecification. We provide an algorithm for estimation using the ML method along with the EM algorithm. Next, we perform comparisons of designs and tests based on statistical power. In addition, we investigate the case of having a large base data set under different scenarios to see how an optimal design can be gained. Finally, we point out current shortcomings and possible future directions.

## 2 Material and Methods

### 2.1 General genetic model

The derivation proceeds from a multinomial logistic regression approach predicting an individual’s genotype G conditional on trait status (affected or unaffected). We treat bases as a mixture of cases and controls according to the population prevalence K. We eliminate correlation between the parameter of interest and other parameters by subtracting the mean from the independent variable. Indexing trait status by *i* (case: *i* = 1, control: *i* = 0) and genotype status by *g* (*g* = 0, 1, 2 depending on reference allele count), the multinomial logistic model is defined by a logistic function:

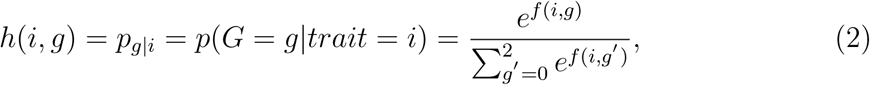

with corresponding *logit* (link) function:

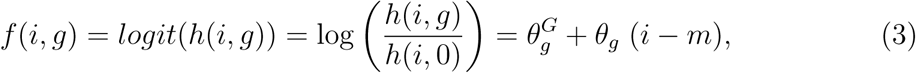

where 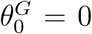 and *m* is the mean value of *i* over all individuals in the dataset under consideration. *θ*_1_ and *θ*_2_ are the log odds ratios for genotypes *G* = 1 and *G* = 2 respectively, relative to the baseline genotype *G* = 0. Under the null hypothesis of interest here (when both *θ*_1_ and *θ*_2_ are zero), 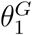 and 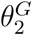 are the log ratios of the frequencies of genotypes *G* = 1 and *G* = 2 relative to the baseline genotype *G* = 0.

### 2.2 Multiplicative trend model

A special case of the general model that is of a particular interest is the multiplicative trend model where *θ*_2_ = 2*θ*_1_ [6]. The *logit* link function can be written as:

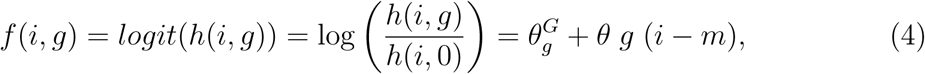

*θ* is equivalent to *θ*_1_ in the general model. Because of its widespread use in genetic epidemiology, we will focus on it for the remainder of this work.

The data for each SNP can be represented as a set of observed genotype counts stratified by sample set (table 1). The conditional probabilities of observing an individual with genotype G=g, given the sample set he/she belongs to, are given in table 2. In the next section, we derive the ST statistic for a univariate analysis of genome-wide SNPs data.

**Table 1:**
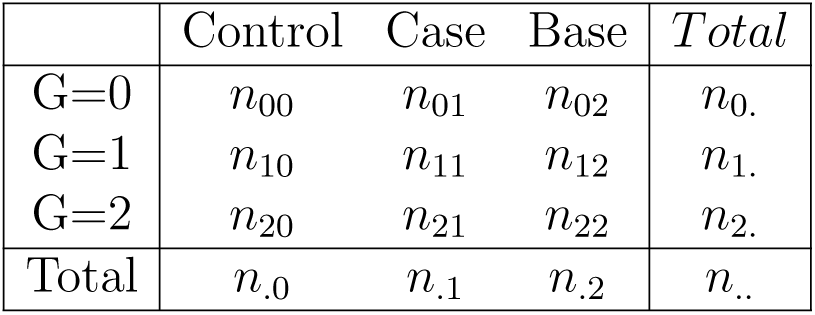
Observed genotype counts, stratified by sample set.

**Table 2:** Probability of genotype conditional on sample set.

|  | Control | Case | Base |
| --- | --- | --- | --- |
| G=0 | $h(0, 0)$ | $h(1, 0)$ | $Kh(1, 0) + (1 - K)h(0, 0)$ |
| G=1 | $h(0, 1)$ | $h(1, 1)$ | $Kh(1, 1) + (1 - K)h(0, 1)$ |
| G=2 | $h(0, 2)$ | $h(1, 2)$ | $Kh(1, 2) + (1 - K)h(0, 2)$ |

### 2.3 Score test for association

The test is derived using the derivative of the likelihood with respect to the parameter of interest (here the log odds ratio), with other parameters (here intercepts) set to their null values. The ST is therefore computationally efficient in the sense that it allows a large number of tests to be performed in a relatively short time compared to other tests. For example, Wald’s test requires the standard error (SE) of the parameter of interest to be evaluated under the alternative hypothesis, and the likelihood ratio test requires estimation of parameters under both the null and the alternative hypotheses. Accordingly, in terms of computational speed they are not as efficient for a genome-wide scan as the ST.

In the CBC design, the conditional counts are assumed to be coming from three independent multinomial distributions; i.e., (*n*_0*i*_, *n*_1*i*_, *n*_2*i*_) ∼ *multinomial*(*n*_.*i*_, *h*(*i*, 0), *h*(*i*, 1), *h*(*i*, 2)), the log-likelihood function using tables 1 and 2 is given by:

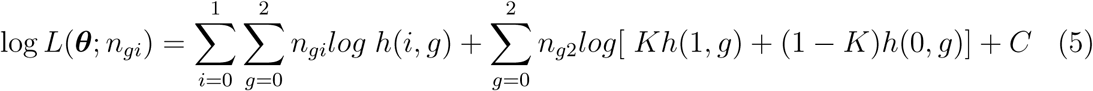

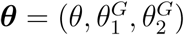. Here we consider a hypothesis test of the multiplicative trend model of the form:

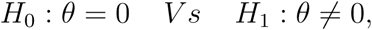

where *θ* is the log odds ratio parameter. We will treat 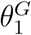 and 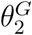 as unknown parameters and K as a known prevalence. The score function is the derivative of the log-likelihood evaluated at *θ* = 0, for this we need the MLE of 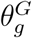 under *H*_0_ which is given by:

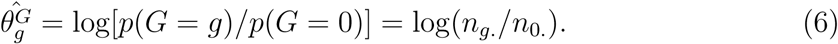

The resulted score function is:

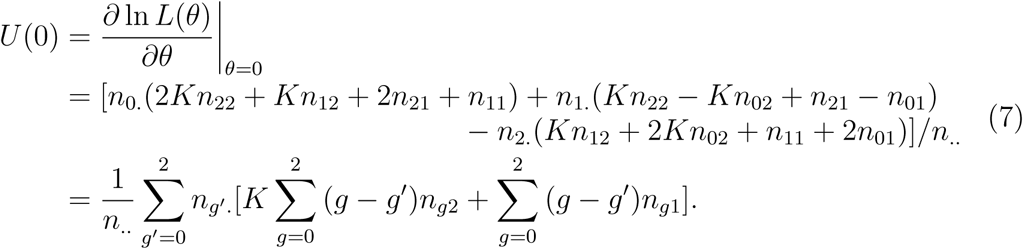

Next, we need the variance of the score function evaluated at *θ* = 0. Since the variance is linear in *n*_*gi*_, we can directly replace the observed cell counts (*n*_*gi*_) by their expected values under *H*_0_ (*n*_.*i*_*n*_*g.*_*/n*_.._). This leads,

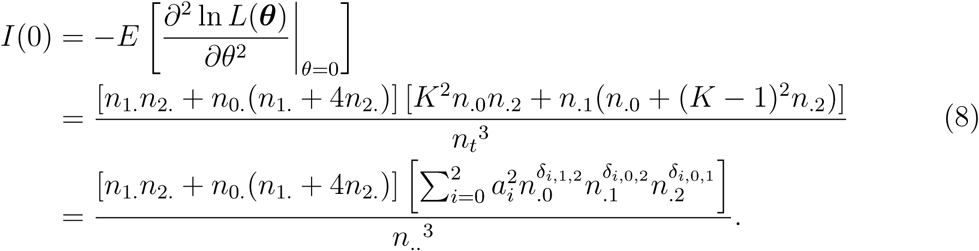

Therefore, the ST statistic is:

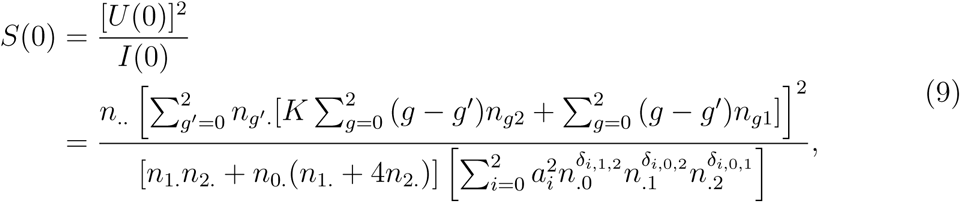

where *a*_*i*_ = 1 −*k, k* and 1 for *i* = 0, 1 and 2. Also *δ*_*i,j,k*_ = 1 if *i* = *j* or *k* and 0 otherwise. Under *H*_0_, this statistic asymptotically follows the central chi-squared distribution with a single degree of freedom 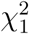. Whereas under *H*_1_ it follows the non-central chi-squared distribution with a single degree of freedom and non-centrality parameter *λ* obtained by replacing observed counts in the ST statistic, by expected cell counts 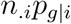 for each of the three groups, using table 2.

It is known that the Cochran-armitage (CA) test is the ST that corresponds to the logistic regression model [7, 8]. Therefore, our ST from the CBC design can be seen as a generalization of the CA test statistic.

### 2.4 Special cases

Here we show the special cases under which the ST from the CBC design is equivalent to the CA test statistic.

- When K=0, the CBC ST statistic reduces to,

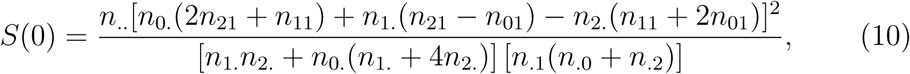

which is the CA statistic, replacing *n*_.0_ by *n*_.0_ + *n*_.2_.

- When K=1, the CBC ST statistic reduces to,

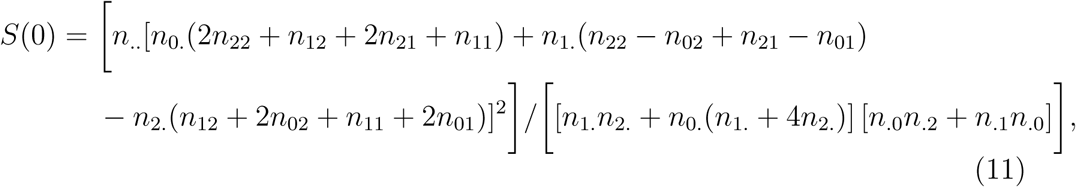

which is also the CA statistic, replacing *n*_.1_ by *n*_.1_+*n*_.2_. The above shows the intertwined between a prospective and a retrospective CBC design. In particular, the special cases of the CBC ST statistic, which were derived using a retrospective regression model are equal to the CA statistic which was derived using a prospective model (supplementary materials). Accordingly, our retrospective approach, which, as described earlier, was merely used for practical reasons is in fact equivalent to the prospective approach, at least when K=0,1. For 0 < *K* < 1, a discussion is given in the supplementary materials.

### 2.5 Score test as a contrast test

The CA test statistic can be seen as a two-sample test for differences between two means; the average number of the reference alleles occurring in the genotypes of cases and that for controls. In contrast, it is unclear which groups’ means are being contrasted under the CBC’s ST given in equation (9). Here, we use simple linear algebra to rewrite the ST in an interpretable way. Specifically, we show that the ST derived from the CBC design can be seen as a contrast test of the form *H*_0_ : *L* = 0 versus *H*_1_ : *L* ≠ 0 where *L* = *n*_.1_(*n*_.0_ + *n*_.2_)(*G*_1_ − *G*_0,2_) + *Kn*_.2_(*n*_.0_ + *n*_.1_)(*G*_2_ − *G*_0,1_) is a contrast of 4 means (*G*_1_, *G*_2_, *G*_0,1_, *G*_0,2_),

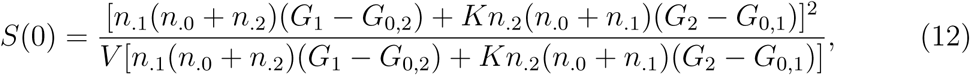

where,

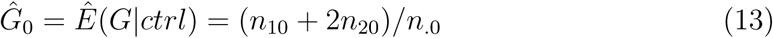

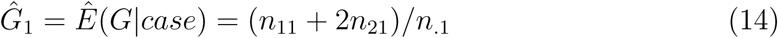

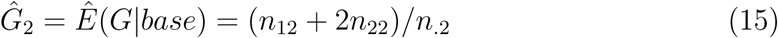

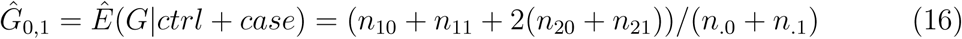

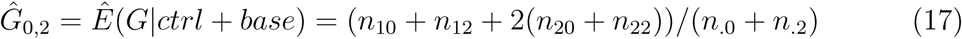

The interpretation of this result goes beyond what the CBC likelihood equation shows. The likelihood which is the joint distribution of the counts across the three groups, shows only the counts information. The score statistic on the other hand, test whether there is a statistically significant deference between these counts. A typical genetic association testing compares the allele frequency mean in two groups; cases and controls. However, since the CBC likelihood allows for information from a third group that is the base set, its score test will not be a simple test of the difference between two means. Indeed, the demonstration above showed how the CBC score test appeared to be contrasting 4 means. This contrast allows for the comparison of two average means; the average of allele means in cases and bases with that in the merged group of controls and bases and merged groups of controls and cases (proof is given in the supplementary materials).

### 2.6 ML estimation

In constructing the ST, we used constrained MLEs (i.e., estimates derived under the null only). Herein, we use the ML method to estimate the underlying parameters, in general, to facilitate the construction of other association tests such as Wald’s and likelihood ratio tests. The derivatives of the log-likelihood function in equation (5) with respect to 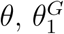 and 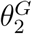 are given by,

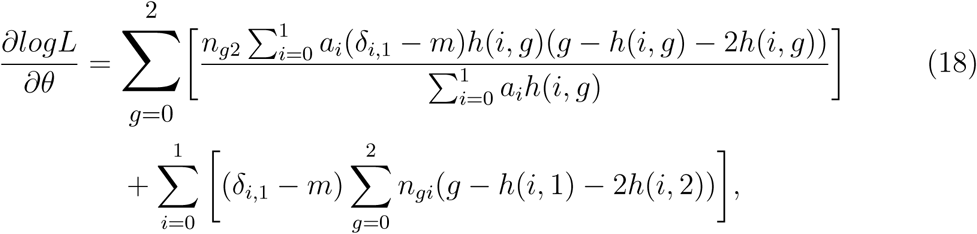

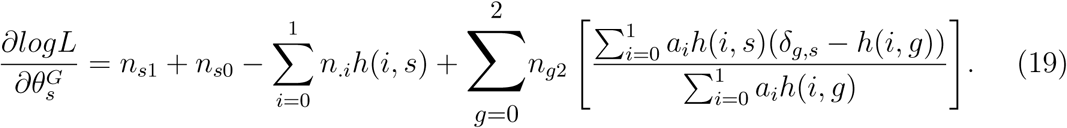

Equating the right-hand sides to zero and solving does not yield a closed form analytical solution. Therefore, a numerical solution is needed. The EM algorithm is used here to overcome convergence problems that arise from having singular Jacobian matrices. Closed forms of the second derivatives to form the Fisher-information matrix are given in the supplementary note.

To illustrate the EM algorithm, we start by treating the counts as an incomplete data. For this, we think of the bases as a multinomial experiment with 6 categories. The counts are (*n*_020_, *n*_021_, *n*_120_, *n*_121_, *n*_020_, *n*_021_), where *n*_*g*2_ = *n*_*g*21_ + *n*_*g*20_, g=0,1,2 and the corresponding multinomial probabilities are ((1 − *K*)*h*(0, 0), *Kh*(1, 0), (1 − *K*)*h*(0, 1), *Kh*(1, 1), (1 − *K*)*h*(0, 2), *Kh*(1, 2)). The log likelihood function based on the complete data is given by:

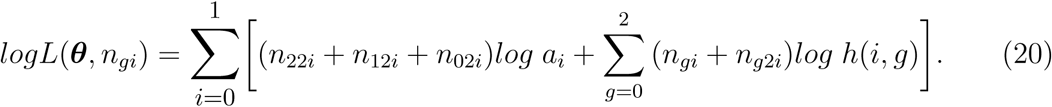

Since *n*_*g*2*i*_ are unobservable, we cannot maximise the above log likelihood directly. This is dealt with using the expectation step. It is known that:

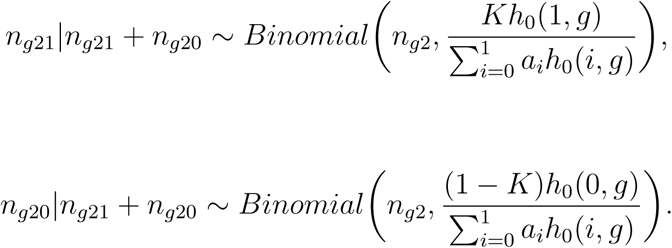

Given an initial guess *θ*_0_ of *θ*, the E-step requires computation of

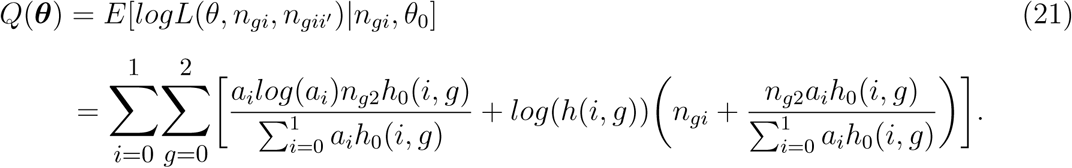

The M-step includes maximising (21) with respect to the parameters to be estimated:

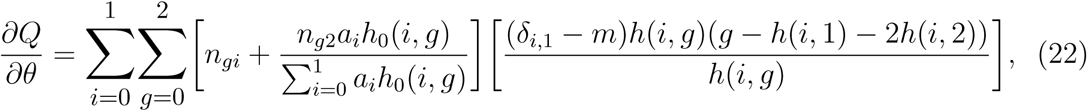

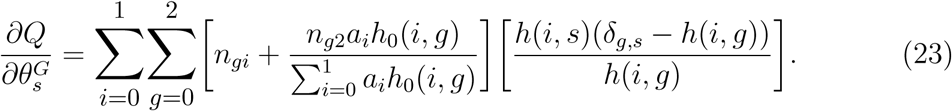

Equating to zero, a numerical solution using either Newton-Raphson or Fisher scoring will be found. In order to implement a practical EM algorithm, two important issues should be taken into consideration. These are the choice of the initials values and the stopping criterion. The choice of the initial values is an important determinant of the number of iterations used in the numerical algorithm to attain the global maximum. A sensible choice is to begin with estimates obtained from a previous GWAS of the same phenotype. Another choice is to use estimates derived from other estimation methods, such as the method of moment. The stopping criterion on the other hand, is equally important and has the potential to make the method less sensitive to the choice of initial values when it is tight enough.

In this paper, all the work is based on simulated data, therefore the parameter values used for data generation were also used as initial values in the estimation process. In a real data application, ideally one would use several initial values over multiple runs of simulations, augmented by a strict stopping criterion, in order to make sure that the algorithm indeed converged to the true global maximum. The stopping criterion we have used is based on the norm of the score function. This norm should be zero when the likelihood attains its global maximum. Therefore, the criterion is that the algorithm stops iterating when the estimated norm of the score function becomes smaller than a pre-specified small constant *ϵ*. The smaller the *ϵ*, the more stringent the criterion is. In the supplementary note, a Mathematica module for estimation is provided to make the task practical.

## 3 Results and discussion

### 3.1 Convergence of score test statistic to asymptotic distribution

As a sanity check, we simulate values of the test statistic, in order to verify its asymptotic distribution under the null and the alternative hypothesis. For this, 5000 small samples (*n*_.1_ = 20, *n*_.2_ = 50 and *n*_.0_ = 30), moderate samples (*n*_.1_ = 200, *n*_.2_ = 500, *n*_.0_ = 300) and large samples (*n*_.1_ = 2000, *n*_.2_ = 5000, *n*_.0_ = 3000) are generated from table 2. The genotype frequencies are taken to be equal to their expected values under Hardy-Weinberg equilibrium, which is the ideal situation; although, this is not required for a genotype-based analysis [9]. Assuming a reference allele frequency p(A)=0.2, the genotype frequencies are *p*_0_ = *p*(*aa*) = 0.64, *p*_1_ = *p*(*Aa*) = 0.32 and *p*_2_ = *p*(*AA*) = 0.04. In all simulation scenarios, K is assumed to be known and equal to 0.2.

Probability-probability plots (pp) provides a heuristic way to check the convergence of the simulated test statistics to the their asymptotic distribution. Figure 1 shows the pp plot of the 5000 simulated test statistics, under the different sample sizes; small, medium and large. The log scale is used to emphasise the small values behaviour (bottom left corner of each plot). It is clear that the deviation from the line y=x over most of the range is small, indicating an appropriate fit between the simulated values and the 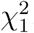. The occasional deviations from the line indicate possible spurious genotype-phenotype associations as well as false negatives, which are both results of departures from the asymptotic chi-squared distribution, due to the finiteness of the samples. Such deviations can be lessened by use of much larger sample sizes, and avoided by use of an exact test, that does not rely on the distribution of the test statistic being correct only asymptotically.

**Figure 1:**
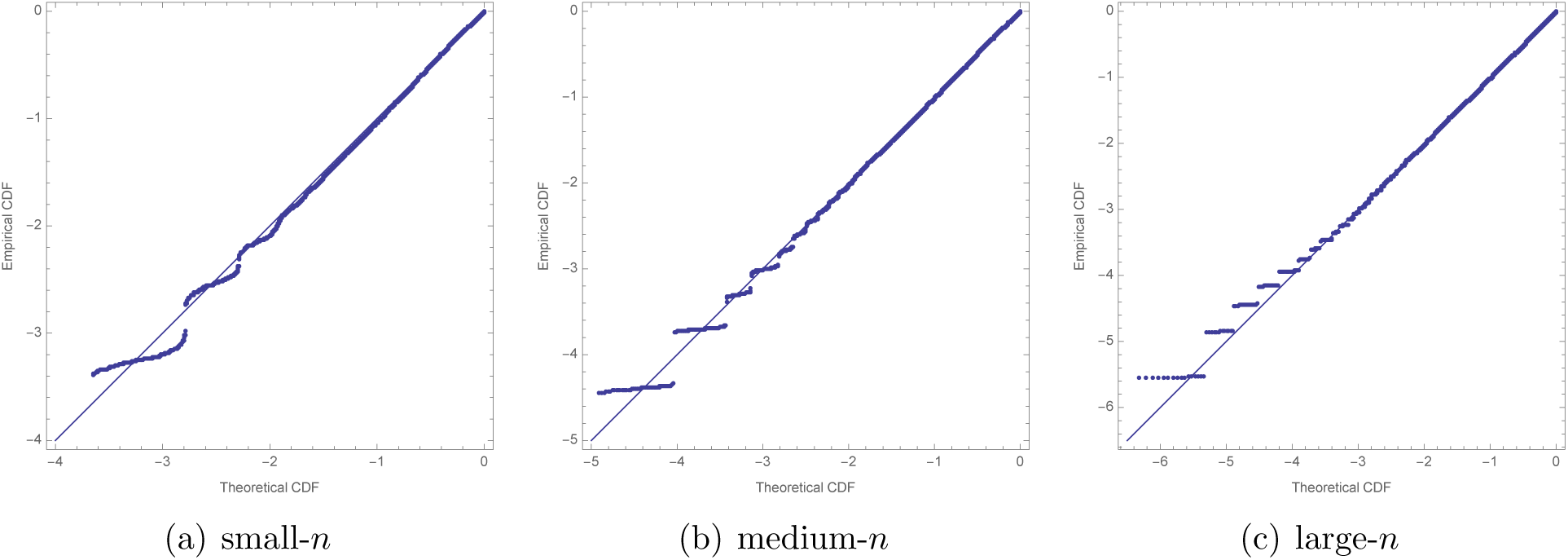
pp plots on log scale of 5000 single-SNP tests of association under the null, assuming K=0.2 is correctly specified. Each plot approximate the blue line, suggesting appropriate adherence to the null distribution 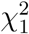. The deviations represent potential spurious associations, and are expected to shrink with larger sample sizes.

Since the prevalence K in our study is assumed to be known, it is constructive to investigate the effect of its misspecification on the asymptotic distribution of the test statistic as well as on statistical power. To this end, we varied the prevalence in the ST statistic and maintained the tables we simulated previously based on a prevalence of 0.2. In Table 3 we summarise the misspecification effect on the distributional behaviour of the simulated test statistics in terms of the area between the probability plot and the straight-line y=x. It is clear that the area becomes smaller as we increase the sample size. Also for medium-*n*, we can see that the smallest area corresponds to the actual value of the prevalence. A similar investigation was carried under the alternative hypothesis with *θ* = *log*[1.5].

**Table 3:** convergence of the ST statistic to its asymptotic distribution under *H*_0_ and *H*_1_ in terms of areas between the pp plot and the y=x line. Actual prevalence is 0.2. The area becomes smaller with increase sample size. Also for medium-*n*, the smallest area corresponds to the actual value of the prevalence.

| | convergence to $\chi^2_1$ under $H_0$ | | | convergence to $\chi^2_\lambda$ under $H_1$ | | |
| --- | --- | --- | --- | --- | --- | --- |
| | small-$n$ | medium-$n$ | large-$n$ | small-$n$ | medium-$n$ | large-$n$ |
| K |  |  |  |  |  |  |
| 0 | 0.007 | 0.0044 | 0.0031 | 0.0145 | 0.0139 | 0.0113 |
| 0.1 | 0.0059 | 0.0038 | 0.0026 | 0.0145 | 0.014 | 0.0107 |
| 0.2 | 0.0071 | 0.0037 | 0.0024 | 0.0136 | 0.013 | 0.0098 |
| 0.3 | 0.0073 | 0.0038 | 0.0031 | 0.0114 | 0.0117 | 0.0083 |
| 0.4 | 0.0093 | 0.0044 | 0.0038 | 0.009 | 0.0094 | 0.006 |
| 0.5 | 0.0073 | 0.0039 | 0.0038 | 0.0055 | 0.0073 | 0.004 |
| 0.6 | 0.0063 | 0.0044 | 0.0045 | 0.0053 | 0.0063 | 0.0023 |
| 0.7 | 0.0055 | 0.0044 | 0.0047 | 0.004 | 0.006 | 0.0018 |
| 0.8 | 0.0075 | 0.0051 | 0.0049 | 0.0061 | 0.0061 | 0.0022 |
| 0.9 | 0.0057 | 0.004 | 0.0041 | 0.0056 | 0.0065 | 0.0031 |
| 1 | 0.0063 | 0.0035 | 0.004 | 0.0067 | 0.0068 | 0.0036 |

The power of the test is defined as the probability of rejecting *H*_0_ when it is false. In other words, it is the probability that the ST statistic will exceed 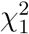, at a significance level *α* of 0.05, under the alternative hypothesis. Accordingly, in our simulation, the power of the ST is the proportion of the simulated test statistics under *H*_1_, which exceed the pre-specified value of 3.84. Figure 2 shows the power as a function of the misspecified prevalence K, using moderate samples. It is encouraging that misspecifying K up to ∼ 0.2 away from the actual value does not lead to a considerable loss of power.

**Figure 2:**
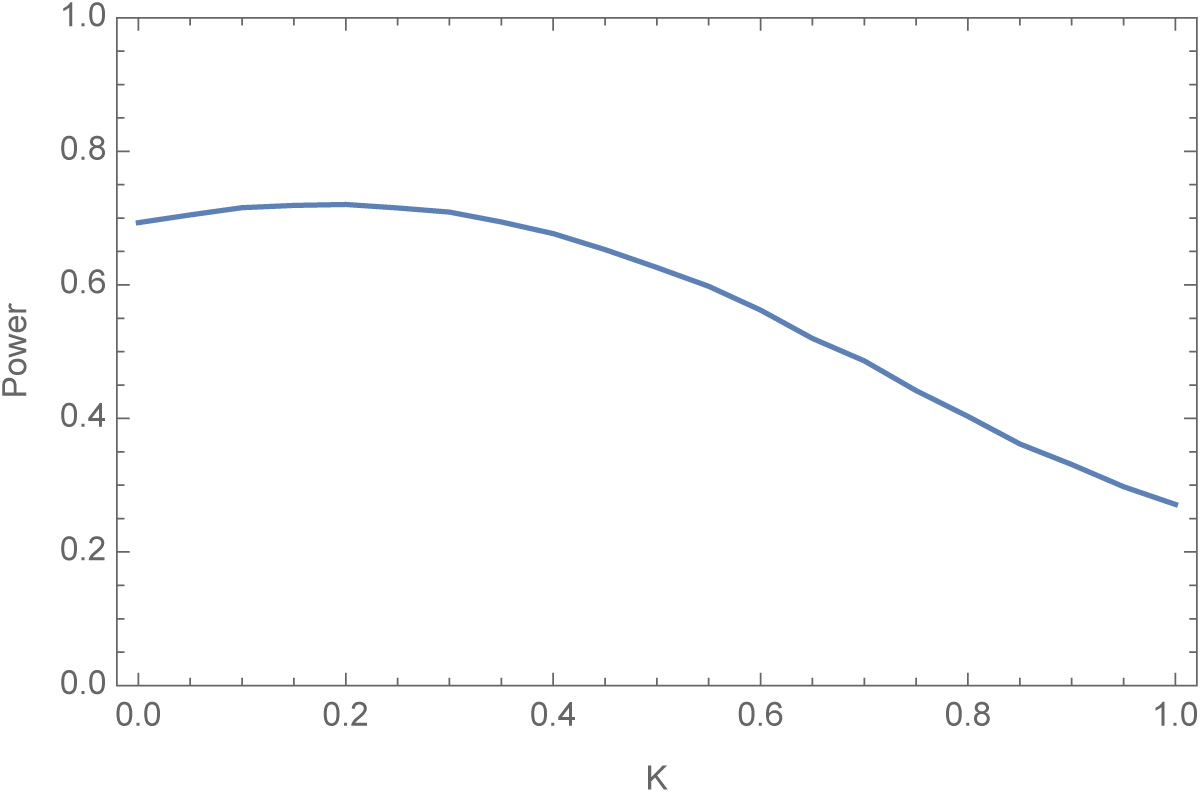
power curve of the score test using medium-*n*: *n*_.1_ = 200, *n*_.2_ = 500 *n*_.0_ = 300 when the actual K=0.2. Misspecifying K up to ∼0.2 away from the actual value does not lead to a considerable loss of power.

### 3.2 Test comparison: LRT, WT and ST

So far we have focused on the ST, merely for practical reasons concerning genome-wide data. Two remaining tests; Wald’s and likelihood ratio, are known to be asymptotically equivalent to the ST. Meaning, their performance in terms of power is approximately the same using large sample sizes. In GWASs, sample sizes are limited, it is therefore constructive to compare their power performance, under moderate sample sizes. To test the null hypothesis of no association *θ* = 0, the WT statistic is given by 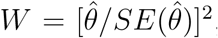, where 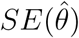 is the non-null SE obtained from the inverse of the Fisher information matrix. On the other hand, the LRT statistic is given by: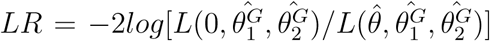, and requires parameter estimation using both the null and the general likelihood.

Using the same simulation procedure under *H*_1_, as before, the empirical power of the three tests is calculated, first under different values of a correctly specified prevalence then assuming a misspecified prevalence. Figure 3 shows the consistency among the three tests, with the LRT showing slightly more power. Note that, when K=0 (1) the CBC design reduced to the usual CC design with number of controls *n*_.0_ + *n*_.2_ (casesn_.1_ + *n*_.2_).

**Figure 3:**
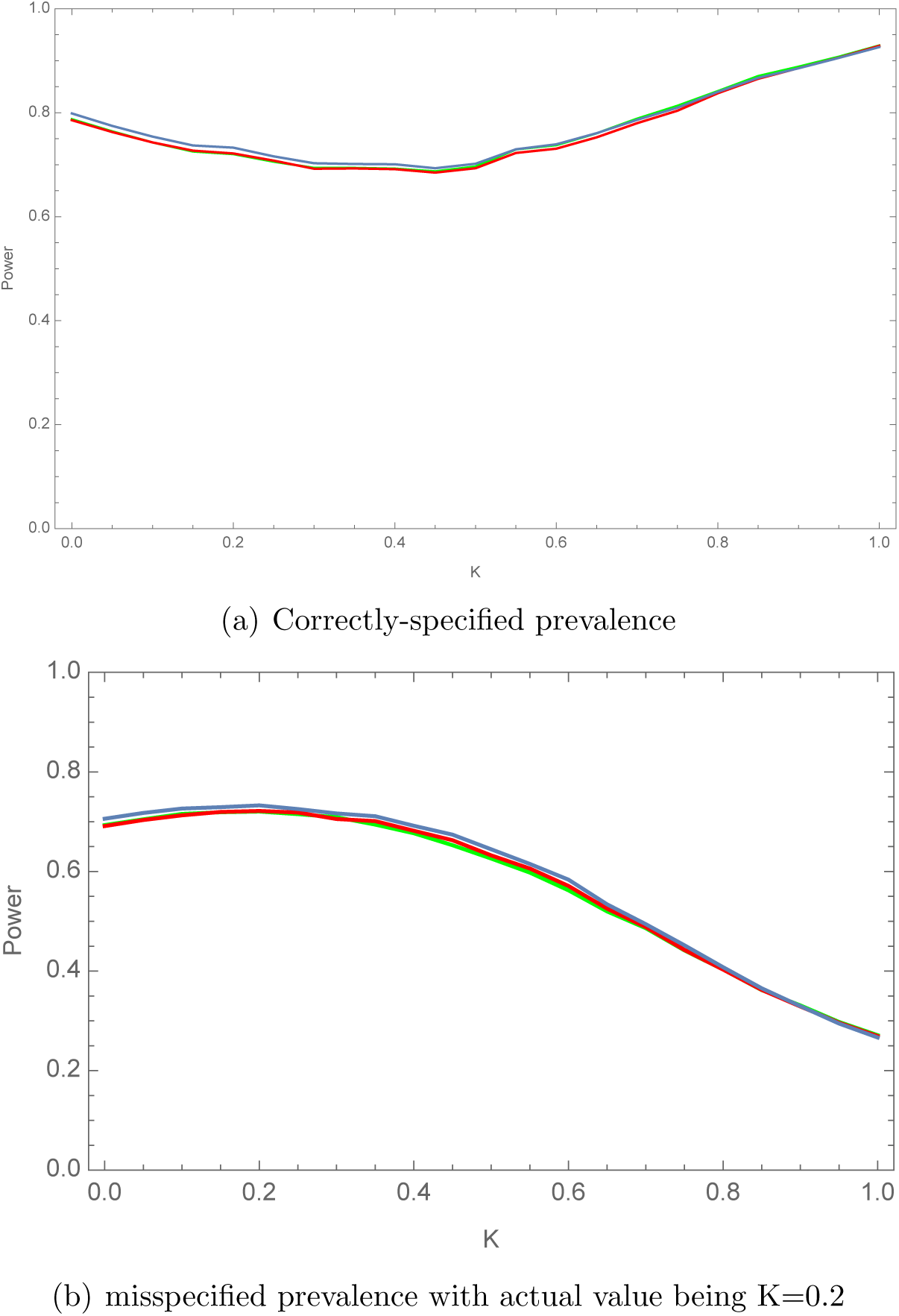
Empirical power of the ST (green curve), WT (red curve), and LRT (blue curve) under the CBC design. Here, medium-*n*: *n*_.1_ = 200, *n*_.2_ = 500 *n*_.0_ = 300 is used.

### 3.3 Design comparison: CC, CB and CBC

Here we compare the ST under the three possible designs; CBC, CC and CB. Figure 4 shows that, the empirical power of the score test under the CBC is equal to that under the CC around K=0.4, which is the proportion of cases in the experiment (cases and controls only). In other words, if *K* ≠ *n*_.1_*/*(*n*_.0_ + *n*_.1_), the CBC design is more powerful than the CC design, which means adding bases is beneficial only if there is imbalance between the proportion of cases in the bases and the proportion of cases in the experiment. As we showed in section 2.4, the ST under the CBC when K=0 (1) is the CA test for a case-control design with 800 controls and 200 cases (700 cases and 300 controls). This explains the higher power at the edges under CBC, compared to the power under the case-control where 200 cases and 300 controls are used. On the other hand, the power of the ST under the case-base design decreases with the prevalence, which is expected because when the trait is common, the case contamination in the bases is high and therefore, treating them as a set of controls can reduce the power.

**Figure 4:**
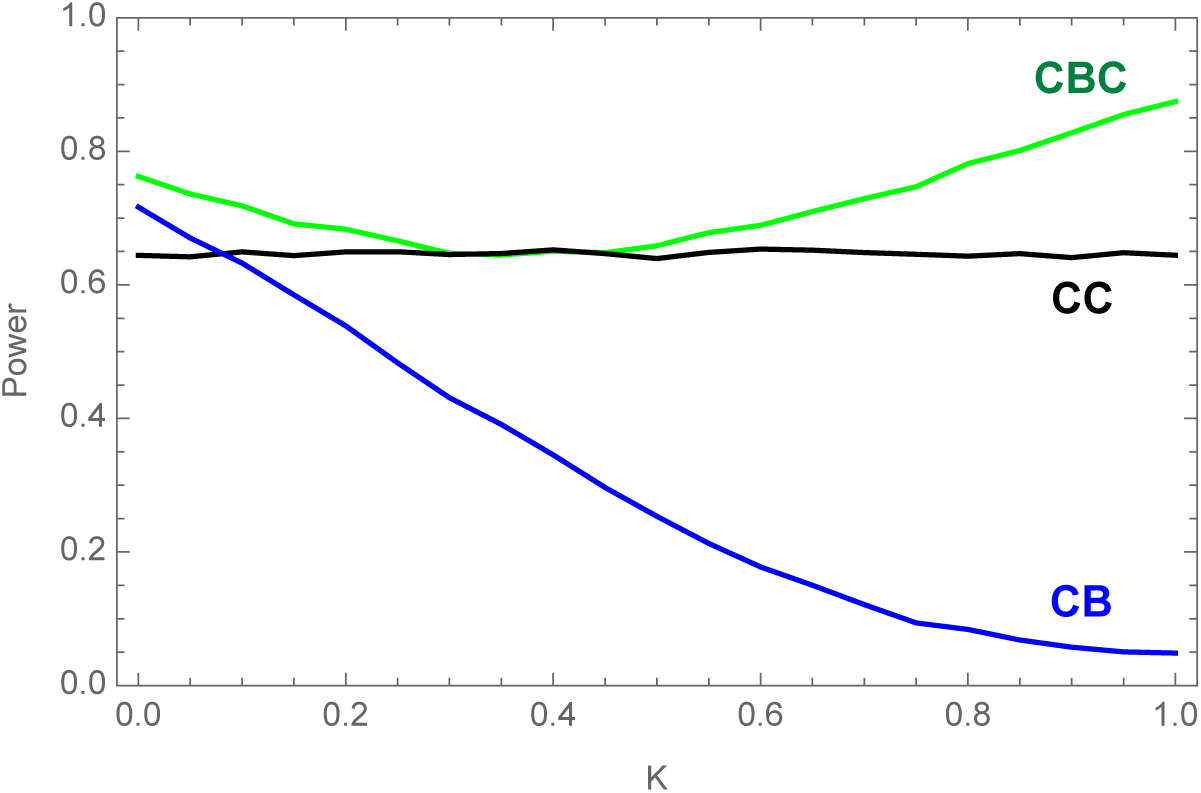
Empirical power of the ST under different designs assuming correctly-specified prevalence. Here, medium-*n*: *n*_.1_ = 200, *n*_.2_ = 500 *n*_.0_ = 300 is used. This means the ST under the CBC, CC and CB uses a sample of size 1000, 500 and 700, respectively. When K=0 (1), the score test from the CBC is the CA test for a case-control design with 800 controls and 200 cases (700 cases and 300 controls). The plot shows (a) the motivation behind the CBC design in the first place, that is the decrease in power under the usual CB design when prevalence increases. (b) The power improvement offered by the CBC design when *K* ≠ *n*_.1_*/*(*n*_.0_ + *n*_.1_).

Above, we assumed that, the CA test can incorporate either controls or bases, but not a combination of them. This is because, in many cases only one of the two sets is available. However, in some cases, where both sets exist, researchers would typically merge bases and controls into one set and analyse it with the cases, using CA test (which is also CBC assuming K=0), which does not take into account any possible misclassification, that is when some of the individuals in the merged set meet the criterion used to define the individuals in the case set. In this case, it will be useful to see whether adding controls to bases would alleviate the loss in power shown above in the CB design, and how this compares to the empirical power of the CBC score test. For this we simulate tables from the CBC design, assuming different values of prevalence. We test for association, first using score test from CBC and then using the above naive approach which treats the combined set of bases and controls as one set of controls. Figure 5, shows this power comparison. Clearly, when K=0, the power of the two tests coincide, as explained in section 2.4. It is also clear that the empirical power of the association test based on the above naive approach of combining bases with controls, still decreases with the prevalence.

**Figure 5:**
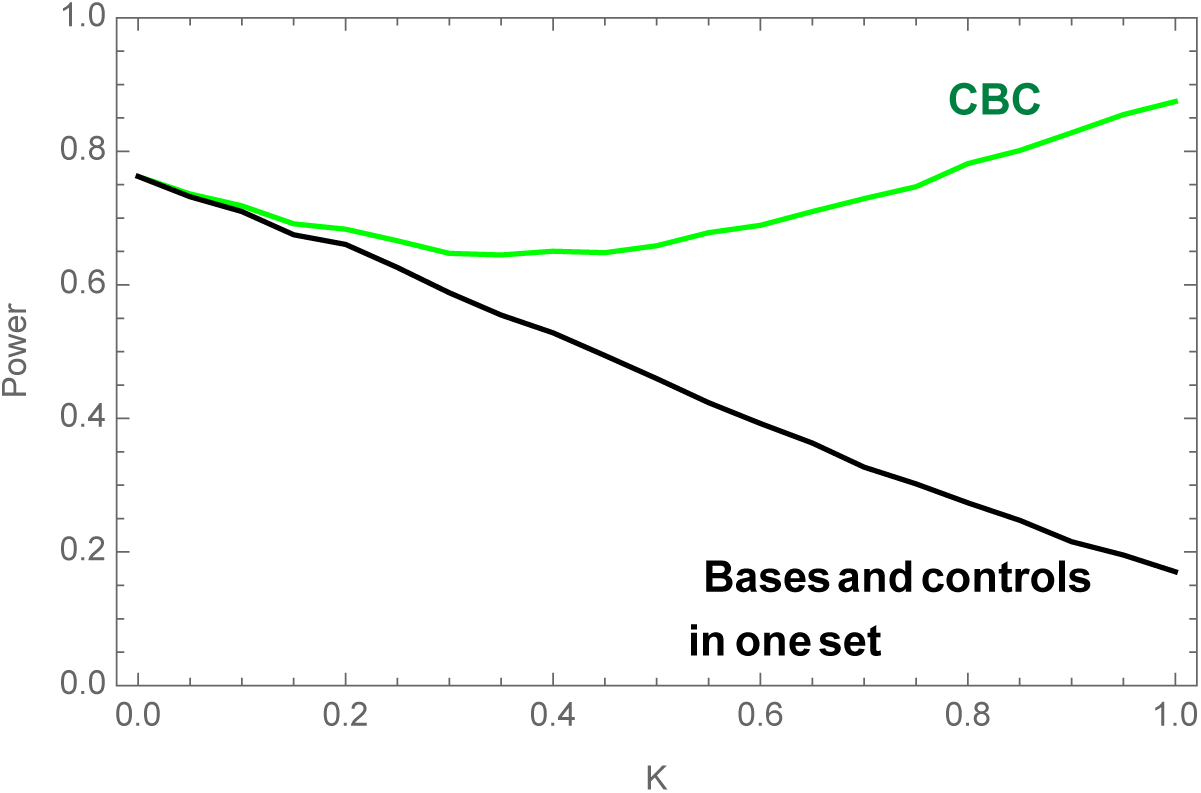
The plot compares the empirical power of the ST from the CBC (assuming correctly-specified prevalence) and that of the CA test after grouping bases with controls. Medium-*n*: *n*_.1_ = 200, *n*_.2_ = 500 *n*_.0_ = 300 was used. This means, both tests use a sample of size 1000. When K≠0, the two tests coincide as shown in section 2.4. When *K* = 0, the empirical power of the CA test when controls are merged with bases, decreases as the prevalence increases. The improvement in power when using the CBC design becomes more evident as K exceeds 0.2

### 3.4 The case of large base data set

The question of whether to collect controls in addition to bases has so far been investigated using a moderate sample size of the base set. Here, we tackle the case of a very large base set e.g. *n*_.2_ = 10000, analysed with a relatively smaller experiment sample *n*_.0_ + *n*_.1_ = 2000.

#### Correctly specified prevalence

Using simulated tables based on an odds ratio of 1.2, figure 6 shows the power as a function of the proportion of cases in the experiment. When K=0.8 (0.2) the power decreases (increases) as the proportion increases, which indicates that an optimal design can be gained by using cases only, when the prevalence is low, and controls only, when the prevalence is high. To confirm this, we fix the number of cases (controls) to be 2000, and plot the power as a function of the number of controls (cases) when K=0.2 (0.8). From figure 7, It is clear that the change in the power as we vary the number of controls (cases) when K=0.2 (0.8) is trivial. In conclusion, an available large base sample set will compensate for missing cases or controls, depending on which of them is more prevalent.

**Figure 6:**
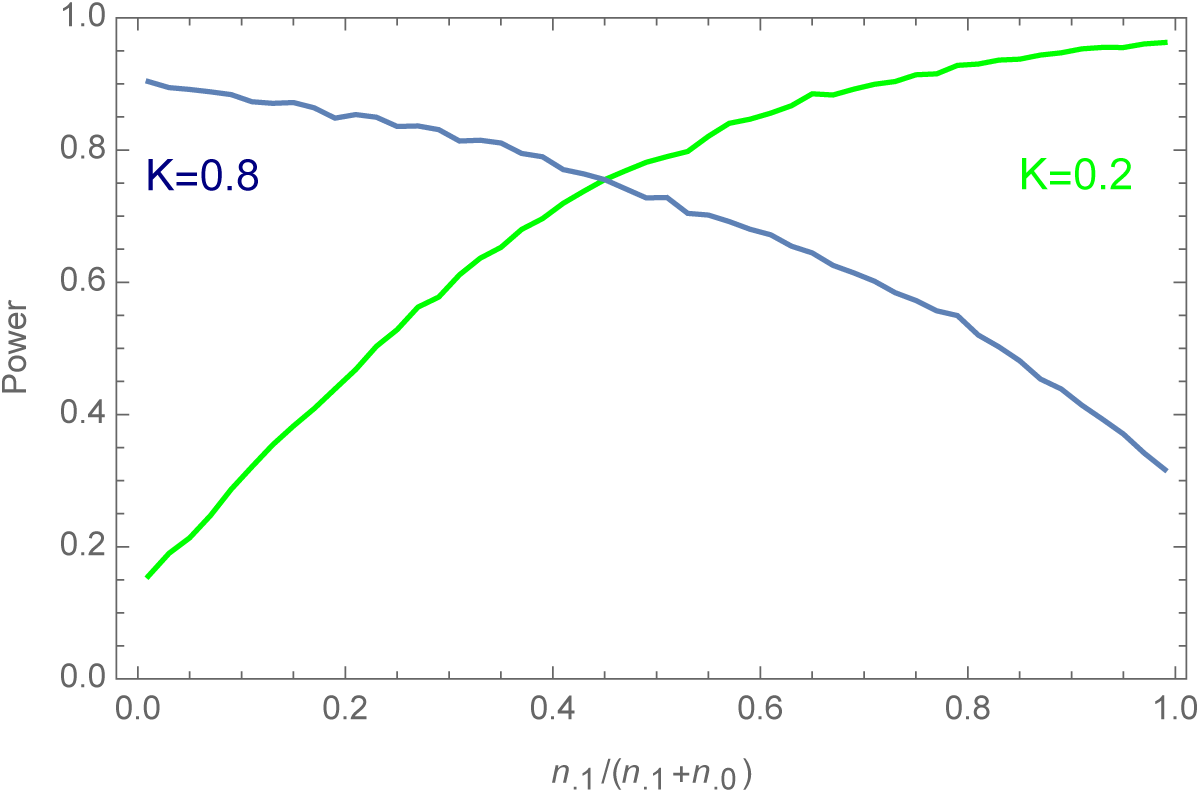
Power as a function of the proportion of cases in the experiment (*n*_.0_ + *n*_.1_ = 2000) using a large set of bases (*n*_.2_ = 10000). The plot suggests that, when the base set is very large, an optimal design can be gained by using cases (controls) only, when the prevalence is low (high).

**Figure 7:**
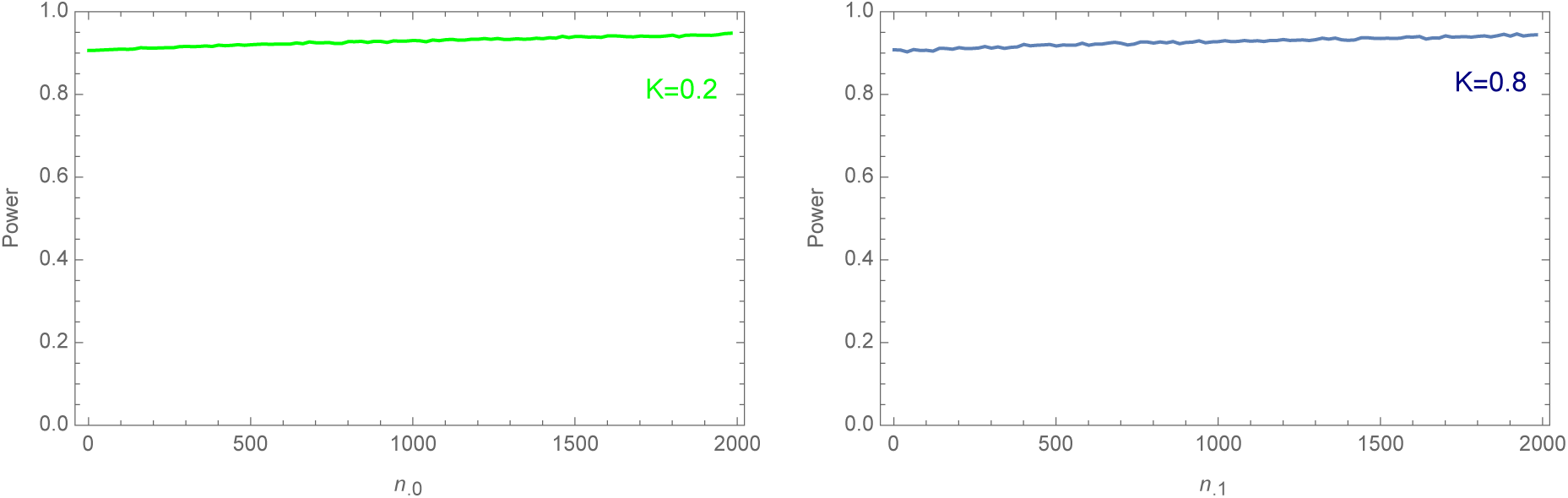
Power as a function of the number of cases (controls) when K=0.8 (0.2) using *n.*0 = 2000 (*n*_.1_ = 2000). There is insignificant change in power when a large base sample set is used.

#### Misspecified prevalence

Assuming only cases have been collected in addition to an available large base sample set, we investigate the same question of whether controls should be collected, under different misspecifications of the prevalence. Figure 8(a) shows the power curve when the actual prevalence is 0.1, 0.3 and 0.5. In each case, we plot the power assuming no misspecification, 10% under and over misspecifications. Power plots show that the test is robust under misspecifications. This is consistent with our observation from figure 2, that misspecifying the prevalence up to 0.2 away from the actual value has a negligible effect on power. Next, we check how serious the misspecification should be to break down the test. Figure 8(b) shows the power curves when the actual value of K is 0.1. The green and blue curves show the power with 20% and 40% over-misspecification. It is clear that still the optimal design can be gained by collecting cases only.

**Figure 8:**
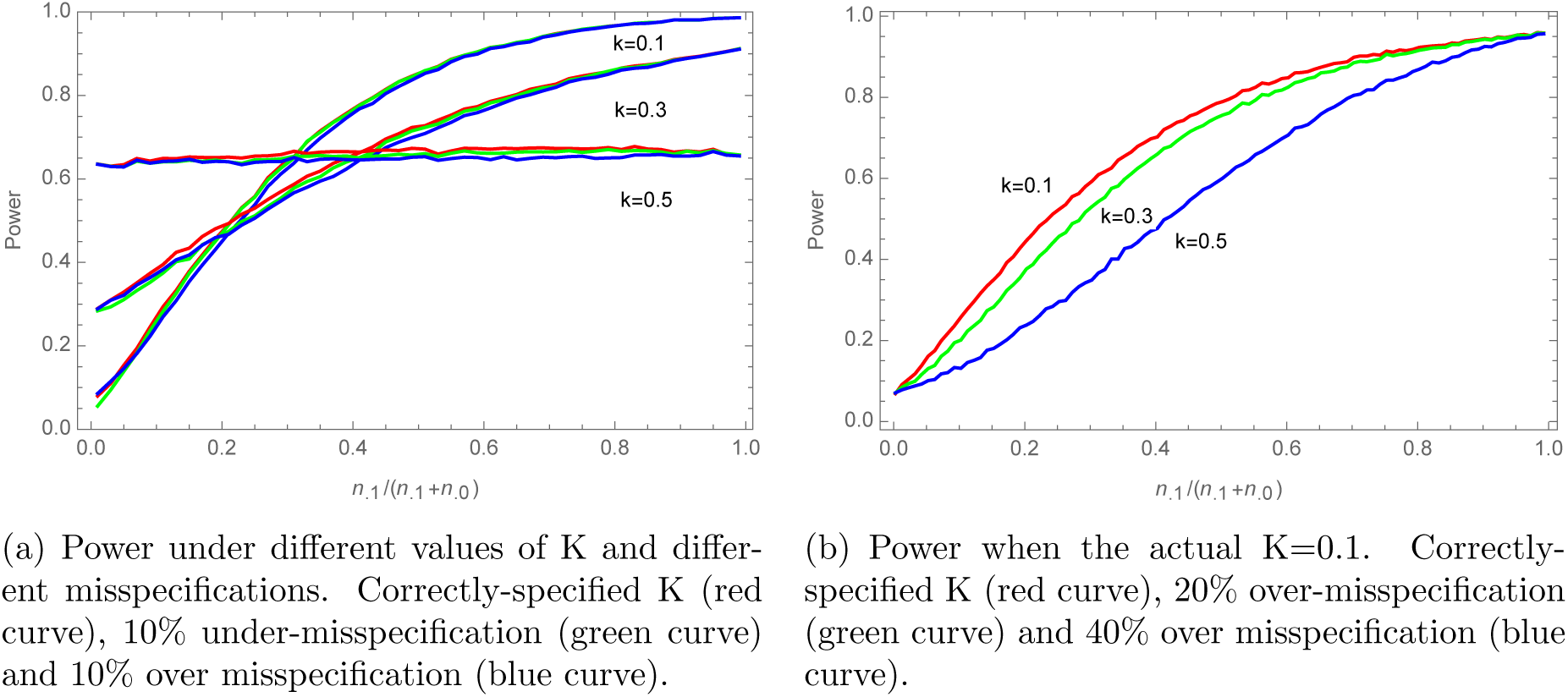
Power under different misspecification using experiment size (*n*_.0_ +*n*_.1_ = 2000) and a large set of bases (*n*_.2_ = 10000). It can be seen that when the base sample set is very large with expected prevalence ≤0.5, the optimal design can be gained by collecting cases only.

## 4 Conclusions

This work has presented a novel CBC method that simulation studies suggest would improve power to identify associations. Application of the CBC to genome-wide data would be useful to illustrate that the method is feasible and can indeed extract new information. One way to show the improvement in power that the CBC offers over other designs, is to establish a pattern of smaller p-values across the genome using the CBC than with other designs. Besides the difficulty of finding large enough studies where such genome-wide pattern of p-values can be observed, the current form of the CBC raises other practical limitations, which make it difficult to perform power assessment, regardless of sample size. The main problems are (a) the difficulty of finding cases, controls and bases genotyped using the same chip. This is essential to avoid assay artefacts; that is the possibility of detecting an association, that is due to differences in chips rather than differences in disease status. (b) The difficulty of finding studies where correction for genetic heterogeneity across the three groups is not essential. This is important in genetic association studies, as even subtle differences between cases and controls, can result in a highly confounded analysis [10].

The current form of the CBC assumes that, disease status is the only covariate in the model. Therefore, in order to allow a swift integration of the data across studies, additional covariates, representing the principal components, should be added to the model to account for differences in genetic backgrounds (population stratification). Similarly, to avoid assay artefacts, a categorical variable should be added to account for batch effects. Finally, although estimates of disease prevalence are often available from previous epidemiological studies, it can be advantageous to estimate it simultaneously with other parameters in the CBC model, by maximizing equation (21) with respect to K too. The module in the supplementary note can be extended accordingly for this purpose.

## 5 ACKNOWLEDGEMENTS

The work described in the paper was funded by the Saudi Government as part of King Saud university scholarships program which the first author was part of. This research project was supported by a grant from the Research Center of the Female Scientific and Medical Colleges, Deanship of Scientific Research, King Saud University. We thank Prof. Ton Coolen for his useful comments.

